# Exploring therapeutic strategies for Infantile Neuronal Axonal Dystrophy (INAD/*PARK14*)

**DOI:** 10.1101/2022.08.16.504080

**Authors:** Guang Lin, Burak Tepe, Geoff McGrane, Regine C. Tipon, Gist Croft, Leena Panwala, Amanda Hope, Agnes J.H. Liang, Zongyuan Zuo, Lily Wang, Hugo J. Bellen

## Abstract

Infantile Neuroaxonal Dystrophy (INAD) is caused by recessive variants in *PLA2G6* and is a lethal pediatric neurodegenerative disorder. Loss of the *Drosophila* homolog of *PLA2G6*, leads to ceramide accumulation, lysosome expansion, and mitochondrial defects. Here, we report that ceramide metabolism, the endolysosomal pathway, and mitochondrial morphology are affected in INAD patient-derived neurons. We show that in INAD mouse models the same features are affected and that glucosylceramides are elevated in dopaminergic neurons and Purkinje cells, arguing that the neuropathological mechanisms are evolutionary conserved and that ceramides can be used as biomarkers. We tested 20 drugs that target these pathways and found that Ambroxol, Desipramine, Azoramide, and Genistein alleviate neurodegenerative phenotypes in INAD flies and INAD patient-derived NPCs. We also develop an AAV-based gene therapy approach that delays neurodegeneration and prolongs lifespan in an INAD mouse model.

**One Sentence Summary:** Ceramide accumulation, lysosomal expansion and mitochondrial defects are a root cause of INAD/PARK14.

## Introduction

Infantile Neuroaxonal Dystrophy (INAD) (OMIM #256600) is a devastating and lethal pediatric neurodegenerative disorder caused by recessive variants in *PLA2G6* (Khateeb et al., 2006; Morgan et al., 2006). Moreover, variants in *PLA2G6* also cause atypical NAD (aNAD) (OMIM # 610217) and *PLA2G6*-related dystonia-parkinsonism, also called Parkinson disease 14 (PARK14) (OMIM #612953) (Paisan-Ruiz et al., 2009). Collectively, these three diseases are called *PLA2G6*-associated neurodegeneration (PLAN) (Kurian et al., 2008). Iron accumulation has been observed in the basal ganglia of some INAD and aNAD patients but not in PARK14 patients. Hence, these diseases are also categorized as neurodegeneration with brain iron accumulation 2 (NBIA2) (OMIM #610217) (Gregory et al., 2008; Morgan et al., 2006). The symptoms of INAD include early onset ataxia (age 1 to 3 years), mental and motor deterioration, hypotonia, progressive spastic tetraparesis, visual impairments, bulbar dysfunction and extrapyramidal signs. INAD patients usually die before their 10^th^ birthday. In contrast to the severe defects in INAD, aNAD and PARK14 patients have a later onset of symptoms, including progressive dystonia and parkinsonism. Cerebellar atrophy is a characteristic symptom in both INAD and aNAD (Sumi-Akamaru et al., 2015). The formation of spheroid structures in the nervous system is a typical neuropathological hallmark of PLAN (Hedley-Whyte et al., 1968). These spheroid structures contain numerous membranes as well as α-synuclein and ubiquitin (Riku et al., 2013) and are named tubulovesicular structures (TVSs) (Sumi-Akamaru et al., 2015). Moreover, Lewy bodies and phosphorylated tau-positive neurofibrillary tangles have also been observed in the nervous system of the PLAN patients (Paisan-Ruiz et al., 2012; Riku et al., 2013).

Mice that lack *PLA2G6* (genotype: *PLA2G6^KO/KO^*) exhibit a normal lifespan but show a slowly progressive motor defect, first observed at around one year of age (Malik et al., 2008; Shinzawa et al., 2008). These mice show some phenotypes observed in INAD patients, including a progressive axonal degeneration (Shinzawa et al., 2008), cerebellar atrophy (Zhao et al., 2011), and the presence of TVS and α-synuclein containing spheroids in the nervous system (Malik et al., 2008; Shinzawa et al., 2008). Transmission electron microscopy (TEM) revealed a swelling of the presynaptic membrane, synaptic degeneration, and mitochondrial inner membrane defects in these mice (Beck et al., 2011).

Another mouse model of INAD that harbors a spontaneous G373R missense mutation in *PLA2G6* (genotype: *PLA2G6^G373R/G37R^*) was also identified. Homozygous *PLA2G6^G373R/G37R^* mice express PLA2G6-G373R protein at a comparable level to wild-type littermates. The inheritance of this mutation is recessive and PLA2G6-G373R protein exhibits no enzymatic activity. Hence, *PLA2G6^G373R/G37R^* is a severe loss of function allele (Wada et al., 2009). Interestingly, these homozygous *PLA2G6^G373R/G37R^* mice only live for about 100 days and hence show severe behavioral and neuropathological phenotypes at a much earlier age than the mice that lack *PLA2G6* (Wada et al., 2009). In addition, no iron accumulation has been documented in the brain of both INAD mouse models.

We previously found that flies that lack *iPLA2-VIA* (the fly homolog of *PLA2G6*), abbreviated INAD flies, display slow progressive neurodegeneration, including severe bang-sensitivity, motor defects as well as defects in vision (Lin et al., 2018). TEM revealed that INAD flies exhibit swelling of synaptic terminals, the presence of TVSs, the disruption of the mitochondrial inner membranes, and an obvious accumulation of lysosomes. The expression of human *PLA2G6* cDNA fully rescues these defects in flies. These data not only show functional conservation of *PLA2G6* throughout evolution but also suggest that introducing human *PLA2G6* into INAD mice or INAD/PARK14 patients may alleviate the progression of the disease.

We previously discovered that iPLA2-VIA binds to and stabilizes the retromer subunits, VPS26 and VPS35 (Lin et al., 2018). Loss of *iPLA2-VIA* reduces the level of VPS26 and VPS35, and impairs retromer function. This leads to an increase in trafficking to the lysosomes followed by an expansion of lysosomes in size and number. This in turn causes an elevation of ceramide levels, increases membrane stiffness which may further impair retromer function, leading to a negative feed-forward amplification of the defects. Pharmacologic or genetic manipulations that either reduce ceramide levels or activate the retromer robustly suppress the loss of *iPLA2-VIA* associated neurodegenerative phenotypes in flies (Lin et al., 2018). Based on our previous findings, we proposed that impaired retromer-endolysosomal function results in an increase in ceramide which is toxic (Lin et al., 2018; Lin et al., 2019).

The previous data raise several questions. Are the defective pathways we observed in flies similarly affected in INAD patient-derived cells as well as INAD mouse models? Can drugs that target ceramide metabolism and the endolysosomal pathway alleviate neurodegenerative phenotypes in INAD models? Finally, can the introduction of human *PLA2G6* cDNA rescue neurodegenerative phenotypes in INAD patient-derived cells and INAD mouse models? Here we report an elevation of ceramides and impaired endolysosomal trafficking and mitochondrial morphology in INAD patient-derived cells as well as in the INAD mice, suggesting that these pathways are affected in all species tested so far. We also assess 20 drugs in INAD flies and patient-derived cells and identify four drugs that improve lysosomal function and reduce elevated ceramides and suppress neurodegenerative phenotypes suggesting a causal relationship. Lastly, we report the development of a gene therapy approach that alleviates neurodegenerative phenotypes and prolongs lifespan in the INAD mice providing potential therapeutic strategies to treat INAD and PARK14.

## Results

### *PLA2G6* is highly expressed in human iPSCs and NPCs, but not in skin fibroblasts

INAD patient-derived skin fibroblasts have been used in several studies to explore the molecular mechanisms of INAD (Davids et al., 2016; Kinghorn et al., 2015; Sun et al., 2021; Villalon-Garcia et al., 2022). We tested the specificity of three commercially available PLA2G6 antibodies and found one antibody that recognizes PAL2G6 (sc-376563 Santa Cruz; Supplemental Figure 1A-B). This antibody recognizes a band of the proper molecular weight in 293T cells as well as in pluripotent stem cells (iPSCs) and Neural Progenitor Cells (NPCs) derived from a healthy person (Supplemental Figure 1A and Figure 1A). Moreover, this band was absent in 293T cells that express PLA2G6 shRNAs and iPSCs and NPCs derived from an INAD patient with a PLA2G6-R70X variant (29-3; see below) (Supplemental Figure 1B and Figure 1A). These data show that this antibody can specifically recognize endogenous PLA2G6.

**Figure 1:**
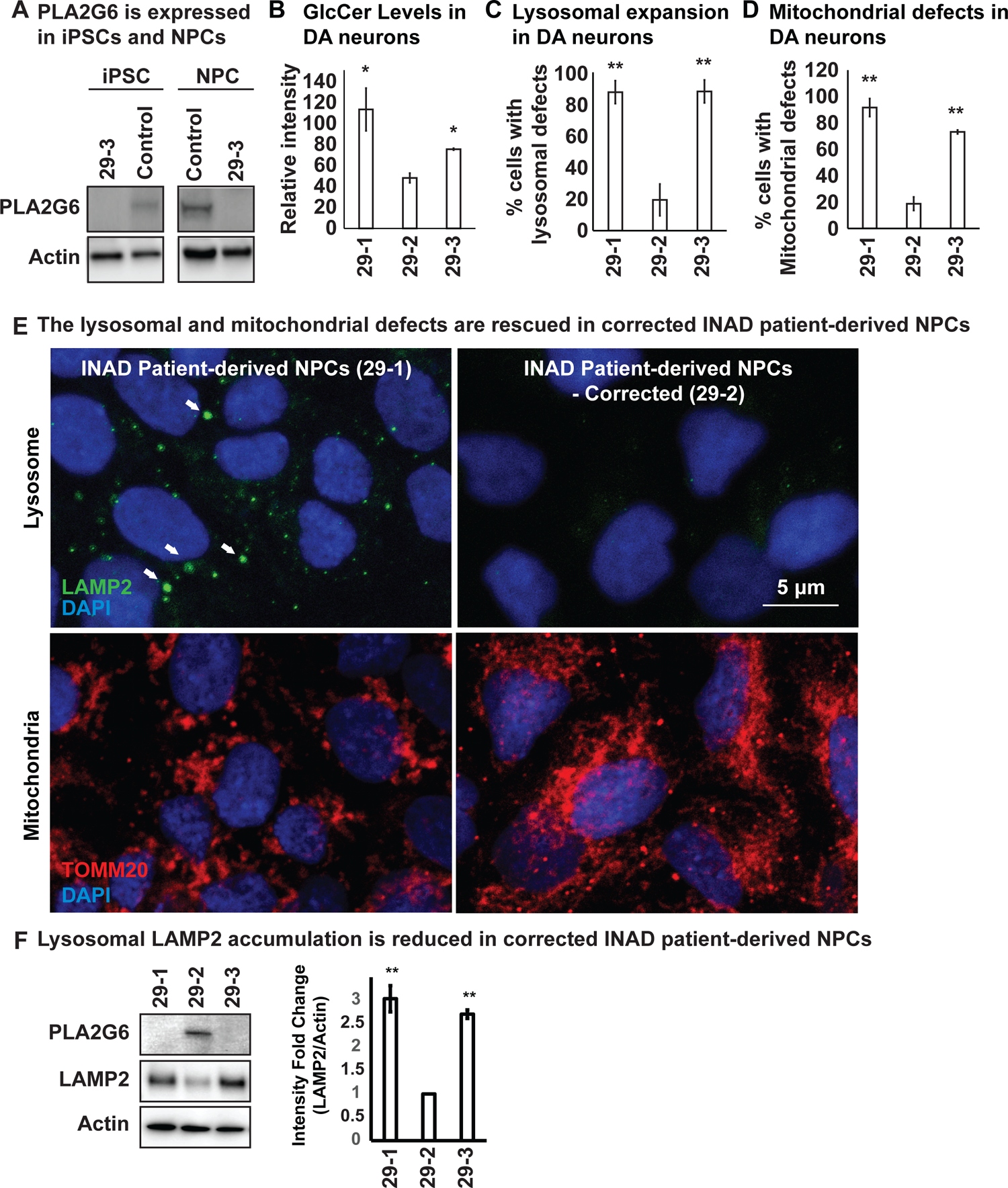
Ceramide accumulation, lysosomal expansion and mitochondrial defects in INAD patient-derived NPCs and DA neurons. A. PLA2G6 is expressed in iPSCs and NPCs using the sc-376563 antibody. The control iPSCs and NPCs were generated by reprograming a fibroblast line from a healthy person (GM23815; Coriell Institute). The 29-3 iPSCs and NPCs were generated from lymphoblasts from an INAD patient in Family 2 (Supplement Figure 1D). Actin was used as a loading control. B. GlcCer levels in DA neurons (images in Supplement Figure 3B) (n=8). C. Lysosomal expansion in DA neurons (Images in Supplement Figure 3B) (n=8-9). D. Cells with mitochondrial defects (Images in Supplement Figure 3B) (n=6-8). Error bars represent SEM; * P<0.05; ** P<0.01. E. The lysosomal and mitochondrial defects are rescued in edited INAD patient-derived NPCs. Immunofluorescence staining of the indicated INAD patient-derived NPCs. LAMP2 antibody (green; arrows) and TOMM20 antibody (red) were used to label lysosomes and mitochondria, respectively. DAPI (blue) labels cell nuclei. Scale bar = 5 µm. F. Lysosomal LAMP2 accumulation is reduced in edited INAD patient-derived NPCs. PLA2G6 antibody was used to detect the endogenous PLA2G6 in the indicated cellular lysates. LAMP2 antibody was used to assess lysosomal accumulation. Actin was used as a loading control. The intensity of the LAMP2/Actin is quantified at the right (n=3). Error bars represent SEM; ** P<0.01.

To explore the expression levels of PLA2G6 in skin fibroblasts, we used skin fibroblasts derived from a healthy person as control. We observed very low expression levels of PLA2G6 (Supplemental Figure 1C). We then obtained skin fibroblasts from two INAD patients and their parents, labeled Family 1 and Family 2 (Supplemental Figure 1D) and observed that all six skin fibroblast lines, including four from unaffected parents and two from INAD patients, express no or very low levels of PLA2G6 (Supplemental Figure 1E).

We previously showed that knocking down *PLA2G6* in Neuro-2A cells leads to the expansion of lysosomes. Moreover, loss of *iPLA2-VIA* in flies leads to the disruption of mitochondrial morphology (Lin et al., 2018). To assess the phenotypes associated with the patient-derived skin fibroblasts, we determined the levels of LAMP2, a lysosomal marker, as well as the morphology of mitochondria. The patient-derived skin fibroblasts (88101) from Family 1 show an elevation of LAMP2 levels and a disrupted mitochondrial morphology (Supplemental Figure 1E-F). However, the patient-derived skin fibroblasts (1914560) from Family 2 do not show any of the phenotypes mentioned above (Supplemental Figure 1E-F). Hence, the phenotypes are inconsistent in patient-derived skin fibroblasts. In summary, skin fibroblasts express no or very low levels of PLA2G6 and exhibit highly variable phenotypes making it difficult to interpret data derived from skin fibroblasts. We, therefore, opted to use iPSCs and neurons derived from these cells to characterize phenotypes associated with INAD.

### Ceramide accumulation, lysosomal expansion, and mitochondrial defects in INAD patient-derived NPCs and DA neurons

To assess the phenotypes associated with INAD patient-derived cells, we reprogramed lymphoblasts from a patient in Family 2 (*PLA2G6-R70X*) into iPSCs. We obtained two clones, 29-1 and 29-3. As mentioned above, 29-3 was used in Figure 1A to determine the expression of PLA2G6. To generate a control for the INAD patient-derived iPSCs, we used CRISPR technology to correct the variant in patient-derived iPSCs (29-1) to generate 29-2 (Supplemental Figure 2A). Hence, 29-1 and 29-2 are an isogenic pair of INAD patient-derived iPSCs. Patient and isogenic gene-corrected control lines were differentiated into ventral midbrain dopaminergic neurons. Each line uniformly expressed the positional transcription factor signature markers for ventral midbrain floor plate region neural progenitor cells (NPC) (OTX2+, LMX1A+, FOXA2+, NESTIN+, and are negative for the forebrain marker FOXG1 (Supplement Figure 2 B and C)(Nolbrant et al., 2017). Upon further differentiation, these NPCs gave rise to a robust population of ventral midbrain dopaminergic neurons (TH^+^) with typical neuronal morphologies (Supplement Figure 2D). We then used the isogenic pair (29-1 and 29-2) of INAD patient-derived ventral midbrain NPCs as well as Dopaminergic neurons (DA neurons) differentiated from these NPCs in subsequent experiments.

**Figure 2:**
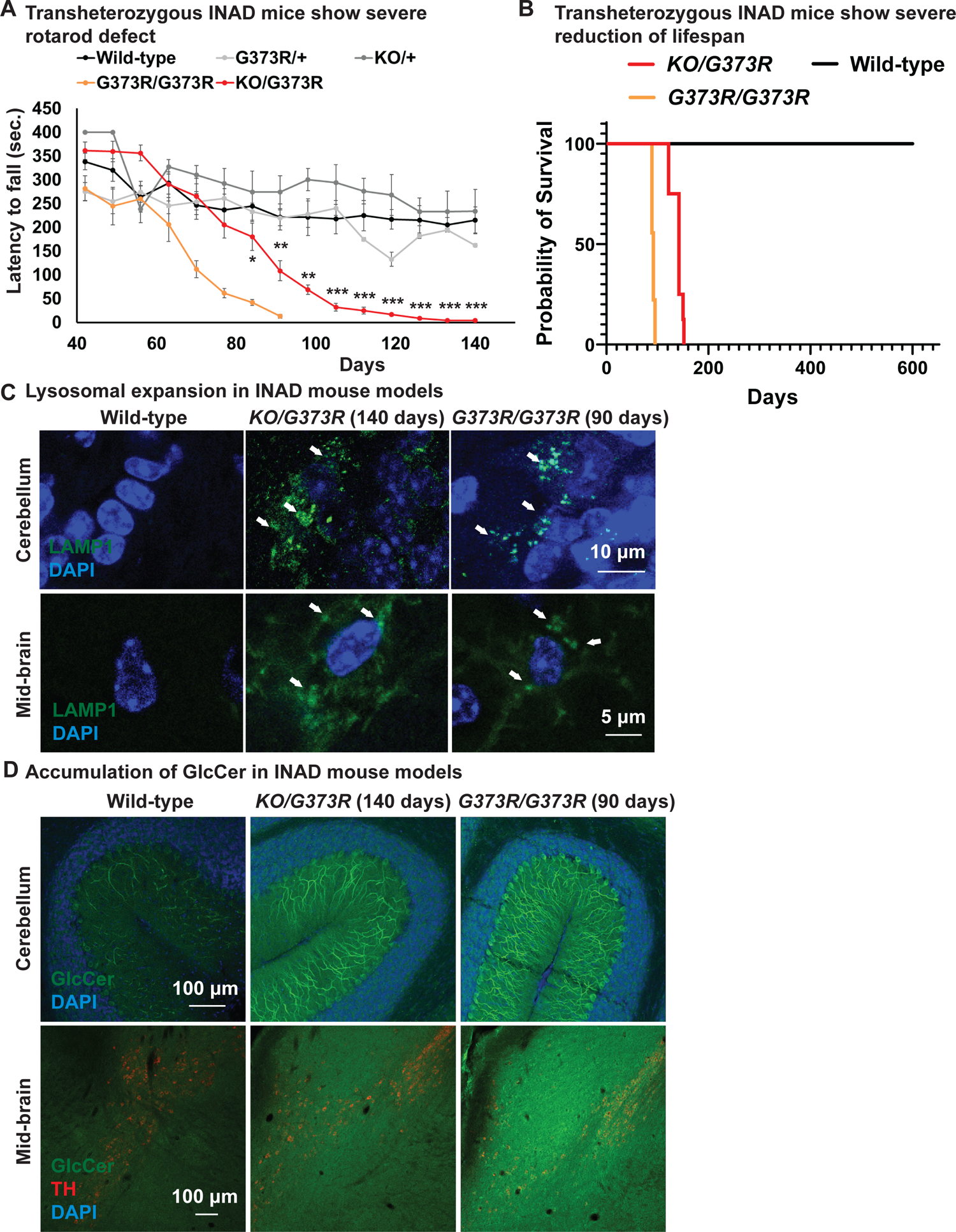
Ceramide accumulation, lysosomal expansion and mitochondrial defects in mouse models of INAD. A. Both *PLA2G6^G373R/G373R^ and PLA2G6^KO/G373R^* mice show severe rotarod defect. Rotarod performance of mice with the indicated genotypes was measured weekly. Wild-type (n=6); +/G373R (n=10); G373R/G373R (n=7); KO/+ (n=4); KO/G373R (n=7). Error bars represent SEM. B. *PLA2G6^G373R/G373R^* and *PLA2G6^KO/G373R^* mice show a severe reduction of lifespan. Wild-type (n=6); *PLA2G6^G373R/G373R^* (n=9); *PLA2G6^KO/G373R^* (n=8). C. Lysosomal expansion in INAD in *PLA2G6^KO/G373R^* and *PLA2G6^G373R/G373R^* mice. Immunofluorescent staining of mouse cerebella and midbrain regions of the indicated genotypes. LAMP1 antibody (green; arrows) labels lysosomes. DAPI (blue) labels nuclei. Scale bar = 10 µm (cerebellum) or 5 µm (midbrain). D. Accumulation of GlcCer in *PLA2G6^KO/G373R^* and *PLA2G6^G373R/G373R^* mice. Immunofluorescent staining of mouse cerebella and midbrain regions of the indicated genotypes. GlcCer antibody (green) was used to assess the levels of GlcCer. TH (Tyrosine Hydroxylase, red) antibody labels DA neurons in the midbrain region. Scale bar = 100 µm. All assays were conducted in blind of genotypes and treatments.

Given that variants in *PLA2G6* also cause parkinsonism, we explored if there are defects associated with INAD patient derived DA neurons. As shown in Supplemental Figure 3A, we did not observe obvious overall cell morphological changes in INAD patient-derived and genetically corrected DA neurons, 29-1 and 29-2, respectively. We measured the levels of Glucosylceramide (GlcCer), and examined the morphology of lysosomes (LAMP2) and mitochondria (ATP5a). GlcCer is easily observed in both control and patient derived DA neurons (Arrows in Supplemental Figure 3B-a and b) but not in the undifferentiated NPCs (Arrowheads in Supplemental Figure 3B-a and b). This suggests that DA neurons, but not NPCs generate GlcCer. However, the GlcCer levels are 2-3 fold higher in INAD patient-derived DA neurons (29-1) than in corrected cells (29-2) (Figure 1B and Supplemental Figure 3B-a and b). Furthermore, lysosomes are increased in number and enlarged in size in cells that carry the variant compared to corrected cells (Figure 1C and Arrows in Supplemental Figure 3B-c and d). Finally, we also observed enlarged mitochondria in the INAD patient-derived DA neurons (29-1) when compared to corrected cells (29-2) (Figure 1D and Supplemental Figure 3B-e to h; 2D).

To extend our observations, we also examined the changes of lysosomal and mitochondrial morphology in NPCs. As shown in Figure 1E, we observed an elevation of the size and number of lysosomes in the INAD patient-derived NPCs (Figure 1E; upper panel). Moreover, the mitochondria in the corrected NPCs form a connected network. However, the mitochondria in the INAD patient-derived NPCs are enlarged and fragmented (Figure 1E; lower panel). To assess the levels of PLA2G6 and LAMP2 we performed western blots on NPCs (Figure 1F; quantified in the right panels). Both clones of the INAD patient-derived NPCs, 29-1 and 29-3, do not express PLA2G6 (Figure 1F; lane 1 and 3, respectively). However, the genetically corrected cells obviously express PLA2G6 (Figure 1F; lane 2). Interestingly, LAMP2 levels are significantly up-regulated in INAD patient-derived NPCs compared to corrected cells (Figure 1F), consistent with a lysosomal expansion. In summary, loss of *PLA2G6* leads to an elevation of GlcCer, expansion of lysosomes, and defects in mitochondrial dynamics. Moreover, these phenotypes are reversed in genetically corrected cells, indicating a causative relationship.

### Ceramide accumulation, lysosomal expansion, and mitochondrial defects in mouse models of INAD

To study the neuropathology in mice, we obtained two INAD mouse models: mice that lack *PLA2G6* (*PLA2G6^KO/KO^*) (Bao et al., 2004) and mice that carry a homozygous *G373R* point mutation (*PLA2G6^G373R/G373R^*) (Wada et al., 2009). These two models exhibit similar neuropathological defects, as explained in the introduction, but the severity of the phenotypes is much stronger in the homozygous *PLA2G6^G373R/G373R^* mice. The *PLA2G6^KO/KO^* mice show a very slow progressive neurodegeneration, whereas the homozygous *PLA2G6^G373R/G373R^* mice display an aggressive and quick neurodegenerative phenotype, yet they were both generated in a *C57BL/6* background. Note that three other *PLA2G6^KO/KO^* have been generated in *C57BL/6* background and they all exhibit very similar very slow progressive phenotypes (Malik et al., 2008; Shinzawa et al., 2008; Zhao et al., 2011). Their lifespan is normal and the first rotarod assay defects are observed at ∼300 days. Moreover, three other models have been described: *PLA2G6^D331Y/D331Y^*, *PLA2G6^R748W/R748W^* as well as a 5’ UTR transposon insertion *PLA2G6^IAP/IAP^* (Chiu et al., 2019; Strokin et al., 2012; Sun et al., 2021). Both point mutation lines, *PLA2G6^D331Y/D331Y^* and *PLA2G6^R748W/R748W^*, express mutant PLA2G6 at a level comparable to their wild type littermates (Chiu et al., 2019; Sun et al., 2021). *PLA2G6^IAP/IAP^* mice produce ∼10% of the wild type protein (Strokin et al., 2012). Given that four mouse models that lack the protein have much less severe phenotypes than the three models that produce PLA2G6 protein, we propose that a compensatory pathway is activated upon a complete loss of *PLA2G6* gene/protein during development. Hence, to minimize genetic background issues and avoid the possible activation of a compensatory pathway we generated and characterized transheterozygous animals (*PLA2G6^KO/G373R^*).

To characterize the *PLA2G6^KO/G373R^* mice, we performed rotarod and lifespan assays. As shown in Figure 2A, the control littermates (*PLA2G6^+/+^* -Wild type; *PLA2G6^KO/+^*; and *PLA2G6^+/G373R^*) show similar normal performances on rotarod assays over a period of 160 days. Homozygous *PLA2G6^G373R/G373R^* mice show the first signs of rotarod defects between 60-70 days of age (Figure 2A) and die within 100 days (Figure 2B). The *PLA2G6^KO/G373R^* mice show the first signs of rotarod defects between 80-90 days of age (Figure 2A) and die at ∼150 days of age (Figure 2B). Hence, we decided to focus on the *PLA2G6^KO/G373R^* mice in the following experiments.

We previously showed that loss of fly homolog of *PLA2G6* leads to an elevation of ceramides including glucosylceramides (GlcCer) (Lin et al., 2018). GlcCer accumulation is also observed in flies that lack the fly homolog of *GBA1* (Wang et al., 2022a), the gene that causes Gaucher disease (Sidransky, 2004; Wong et al., 2004). We performed immunostaining in the cerebellum and midbrain of the *PLA2G6^KO/G373R^* and homozygous *PLA2G6^G373R/G373R^* animals. As shown in Figure 2D, the GlcCer levels are highly up-regulated in the Purkinje cells (Figure 2D; upper panel) and mid-brain cells (Figure 2D; lower panel) of both INAD mouse models when compared to controls, again showing a defective ceramide metabolism.

We next explored the morphology of lysosomes in the cerebellum and midbrain of the *PLA2G6^KO/G373R^* and homozygous *PLA2G6^G373R/G373R^* animals. As shown in Figure 2C, we observed an expansion of LAMP1, a lysosomal marker, in the Purkinje cells in the cerebellum (Figure 2C; upper panel) and neurons of the midbrain region (Figure 2C; lower panel). Hence, lysosomal expansion is a common feature of all models of INAD.

We next performed TEM to assess the morphology of the mitochondria in the cerebellum of the homozygous *PLA2G6^G373R/G373R^* animals. As shown in Supplement Figure 4A, the morphology of mitochondria is disrupted in Purkinje cells of the homozygous *PLA2G6^G373R/G373R^* animals (Supplement Figure 4A). Moreover, we also observed a very significant increase in multivesicular bodies when compared to control animals (Supplement Figure 4B), consistent with an endolysosomal defect. Finally, these mice also exhibit tubulovesicular structures (Supplement Figure 4B), a hallmark of *PLA2G6* mutant animals (Sumi-Akamaru et al., 2015).

In summary, *PLA2G6* mutant mice exhibit a ceramide accumulation, lysosomal enlargement as well as mitochondrial defects. These data are consistent with the fly model of INAD (Lin et al., 2018) and patient-derived NPCs (Figure 1C and D) and DA neurons (Supplement Figure 2B). Hence, we argue that these defects are evolutionary conserved and may play a critical role in the pathogenesis of INAD/PARK14.

### Ambroxol, Azoramide, Desipramine, and Genistein alleviate neurodegenerative phenotypes in INAD flies and patient-derived NPCs

We next tried to identify therapeutic strategies for INAD. We used INAD flies in a primary screen and then tested the drugs that improve the phenotypes in flies in INAD patient-derived cells. Based on a review of the literature, we identified twenty drugs that have been reported to control or affect sphingolipid metabolism, endolysosomal trafficking and drugs that are being tested to treat Parkinson’s disease (Figure 3A; see references in Figure 3A). We previously reported that INAD flies show severe bang-sensitivity at 15 days of age (Lin et al., 2018). This assay is a quick and sensitive assay that measures the propensity of flies to seize upon shock (Figure 3B). Wild-type flies right themselves in less than a second after a vortex paradigm. In contrast, bang-sensitive flies are paralyzed for an extended period of time before they can right themselves. We used this assay as our primary screen assay (Figure 3A). As shown in Figure 3B, wild-type flies are not bang-sensitive. In contrast, INAD flies show severe bang-sensitivity and require ∼20 seconds to recover (Figure 3B). We identified drugs that worsen the bang sensitivity, including Fingolimod, Ozanimod, Fumonisin, Hydrochloroquine, Lonafarnib, Omigapil, and Taurine (Figure 3B). Another set of drugs did not affect bang sensitivity, including Miglustat, Ibiglustat, NCGC607, Rapamycin, VER-155008, Deoxygalactonojirimycin, CuATSM and Metformin (Figure 3B). However, five drugs suppressed bang sensitivity, including Ambroxol, Genistein, PADK, ML-SA1 and Azoramide (Figure 3B). To confirm the rescue effect, we retested the drugs that suppressed the bang sensitivity at a dosage that is 10 times higher than the dosage used in the first round. Each drug exhibited a dose-dependent increase in rescue activity, suggesting that the ability to suppress bang sensitivity is dose-dependent (Figure 3B).

**Figure 3:**
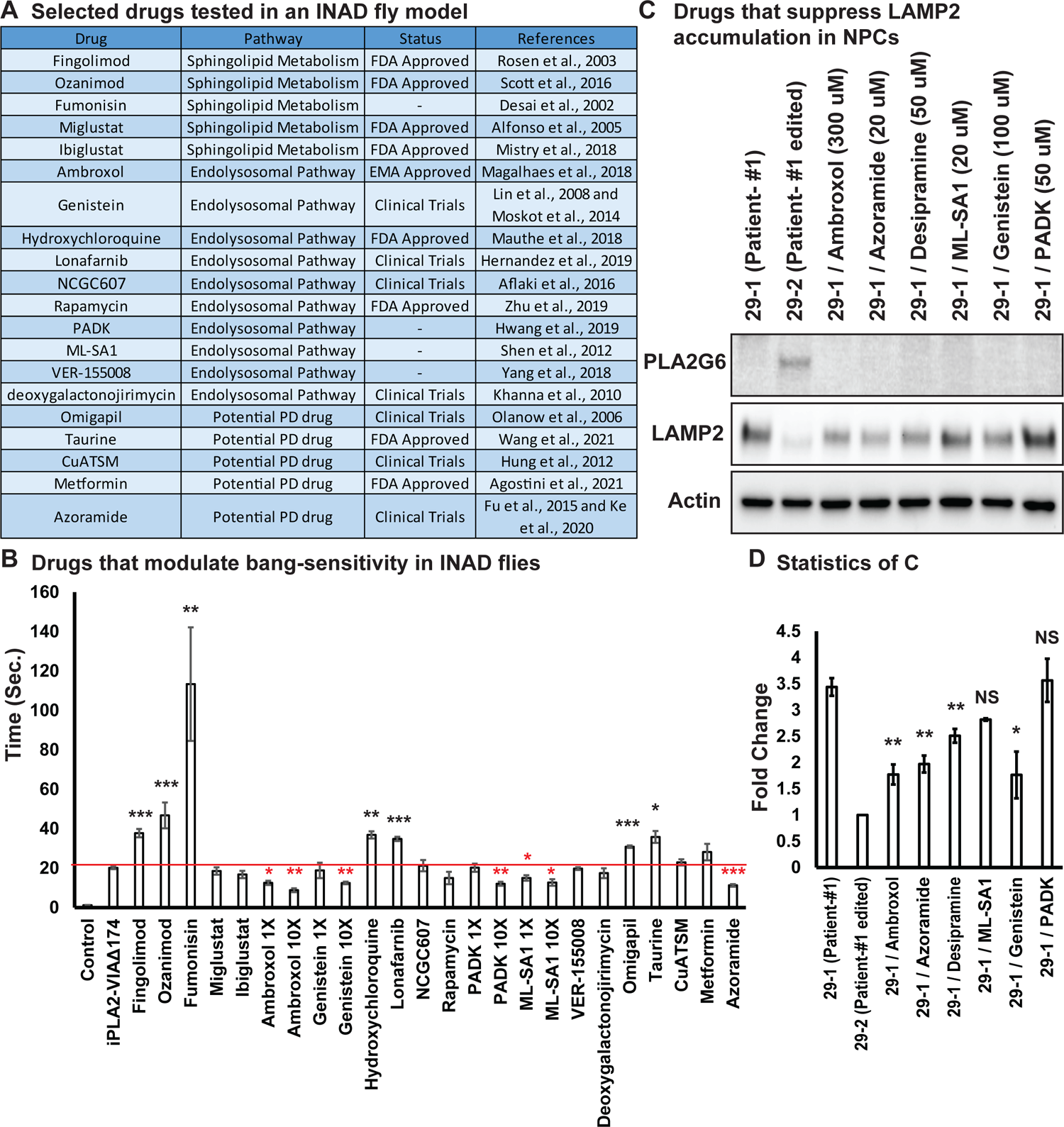
Ambroxol, Azoramide, Desipramine and Genistein alleviate neurodegenerative phenotypes in INAD flies and patient-derived NPCs. A. Selected drugs tested in an INAD fly model. B. Bang sensitivity was used as a primary readout to select drugs that suppress neurodegeneration. Bang sensitivity of control or INAD flies fed with the indicated drugs. Error bars represent SEM (n=3; 20 flies per assay). * P<0.05; **P<0.01; *** P<0.001. Redline highlights the time required for INAD flies to recover from bang-induced paralysis. Red “*” indicates drugs that significantly suppress bang-sensitivity. Black “*” indicates drugs that significantly promote bang-sensitivity. C. Using INAD patient-derived NPCs to select drugs that suppress LAMP2 accumulation. PLA2G6 antibody was used to detect the endogenous PLA2G6 in the indicated cellular lysates. The intensity of the LAMP2/Actin is quantified at D. (n=3). Error bars represent SEM; * P<0.05; ** P<0.01; NS: not significant. References in Figure 3A: (Aflaki et al., 2016; Agostini et al., 2021; Alfonso et al., 2005; Desai et al., 2002; Fu et al., 2015; Hernandez et al., 2019; Hung et al., 2012; Hwang et al., 2019; Ke et al., 2020; Khanna et al., 2010; Liu et al., 2008; Magalhaes et al., 2018; Mauthe et al., 2018; Mistry et al., 2018; Moskot et al., 2014; Olanow et al., 2006; Rosen and Liao, 2003; Scott et al., 2016; Shen et al., 2012; Wang et al., 2021; Yang and Tohda, 2018; Zhu et al., 2019)

To assess if these drugs modify the phenotypes in INAD patient-derived NPCs we probed Western blots of the treated cells for LAMP2. As shown in Figures 3 C and D, patient-derived NPCs (29-1) do not express PLA2G6 and exhibit elevated levels of LAMP2. Correcting the variant (29-2) strongly reduces LAMP2 levels. Importantly, Ambroxol, Azoraminde and Genistein, significantly reduce LAMP2 levels in the patient-derived NPCs (Figures 3C and D).

We previously showed that Myriocin, R55 and Desipramine reduced ceramide levels as well as the lysosomal expansion, and alleviate bang-sensitivity in the INAD fly model (Lin et al., 2018). Given that Myriocin is toxic in vertebratesand that R55 poorly penetrates the blood-brain barrier we tested Desipramine, an FDA approved drug, in the INAD patient-derived NPCs. As shown in Figures 3 C and D, Desipramine also reduces LAMP2 levels in patient-derived NPCs. In summary, we identified four drugs that suppress the loss of *PLA2G6*-induced phenotypes in flies and INAD patient-derived cells: Ambroxol, Azoraminde, Desipramine and Genistein.

### Expression of human *PLA2G6* restores lysosomal and mitochondrial morphology defects in INAD patient-derived NPC lines

We previously showed that whole body expression of human *PLA2G6* in INAD flies fully rescued the neurodegenerative phenotypes and lifespan (Lin et al., 2018). In contrast, neuronal expression of human *PLA2G6* strongly suppressed neurodegenerative phenotypes, but did not prolong lifespan. These data suggest that even though *PLA2G6* plays an important role in the nervous system, it is also required in cells other than neurons. We, therefore, surmised that a gene therapy approach in mice should attempt the delivery of human *PLA2G6* into as many cell types as possible. We, therefore, designed an AAV-based gene therapy construct, *AAV-EF1a-hPLA2G6* (Figure 4A). We used elongation factor EF-1 alpha (EF1a), a ubiquitous promoter, to express *PLA2G6* as broadly as possible. We also designed two constructs, *Lenti-CMV-hPLA2G6,* and *AAV-EF1a-EGFP* as controls (Figure 4A). The lentivirus-based construct allowed quite elevated levels of expression and was used to test toxicity when *PLA2G6* is highly expressed in NPCs. The *AAV-EF1a-EGFP* construct was used to track viral transduction and expression efficiency. Wild-type mice injected with the *AAV-EF1a-EGFP* (serotype) construct via Intracerebroventricular (ICV) and Intravenous (IV) injections at P40 express EGFP in numerous tissues, including the cerebellum, olfactory bulb, cerebral cortex, mid-brain region, spinal cord, sciatic nerve, heart and liver (Supplemental Figure 5). However, EGFP is not expressed in the eyes (Supplemental Figure 5). These data show that *AAV-EF1a-EGFP* is broadly expressed in many tissues when the virus is delivered via ICV and IV. However, not all cells express EGFP based on this assay.

We then conducted a pilot experiment in HEK-293T cells to assess the expression of the protein using these constructs. As shown in Figure 4B, HEK-293T express very low levels of endogenous PLA2G6 (upper bands marked by a *). Note that we also observed a nonspecific band (lower bands marked by #). As expected, *Lenti-CMV-hPLA2G6* induces very high levels of expression of human PLA2G6 in HEK-293T cells (stars in Figure 4B; last lane). However, expression of human PLA2G6 by infecting cells with *AAV-EF1a-hPLA2G6* packaged in either the AAV-PHP.eB or AAV9 serotype is very low, even when we used a very high multiplicity of infection (MOI of 1,000). (Stars in Figure 4B; lanes 3 and 4).

Upon showing that these constructs can be expressed in HEK-293T cells, we tested their expression and function in the patient-derived NPCs (Figure 4C). Expression of PLA2G6 is easily detectable in NPC cells. Moreover, the Lenti-CMV-hPLA2G6 derived protein is expressed at very high levels (∼9 fold higher than the endogenous levels) levels in the NPCs (Figure 4C; last lane). However, expression of human PLA2G6 driven by *AAV-EF1a-hPLA2G6* construct packaged in AAV-PHP.eB or AAV9 serotype is very low (∼10% of the endogenous levels) (Figure 4C; lanes 3 and 4). Interestingly, the LAMP2 levels are reduced by about 10-20% (Figure 4C; bottom) in all three conditions, suggesting that even very low levels may have a beneficial effect.

We also examined mitochondrial morphology in these cells. As shown in Figure 4D, the patient-derived NPCs show enlarged and fragmented mitochondrial morphology (Figure 4D; upper left). AAV-PHP.eB-hPLA2G6 (Figure 4D; upper right), and AAV9-hPLA2G6 (Figure 4D; lower left) strongly rescues this morphological abnormality. In contrast, the Lenti-CMV-hPLA2G6, which expresses very high levels of human PLA2G6, only partially rescues mitochondrial morphological abnormalities (Figure 4D; lower right). This suggests that very high levels of expression of PLA2G6 may be somewhat toxic. In summary, both AAV-PHP.eB-hPLA2G6 and AAV9-hPLA2G6 induce low levels of expression of human PLA2G6 in NPCs (10-20% of the endogenous levels), and can partially alleviate two key phenotypes: lysosomal expansion and mitochondrial morphological abnormalities.

**Figure 4:**
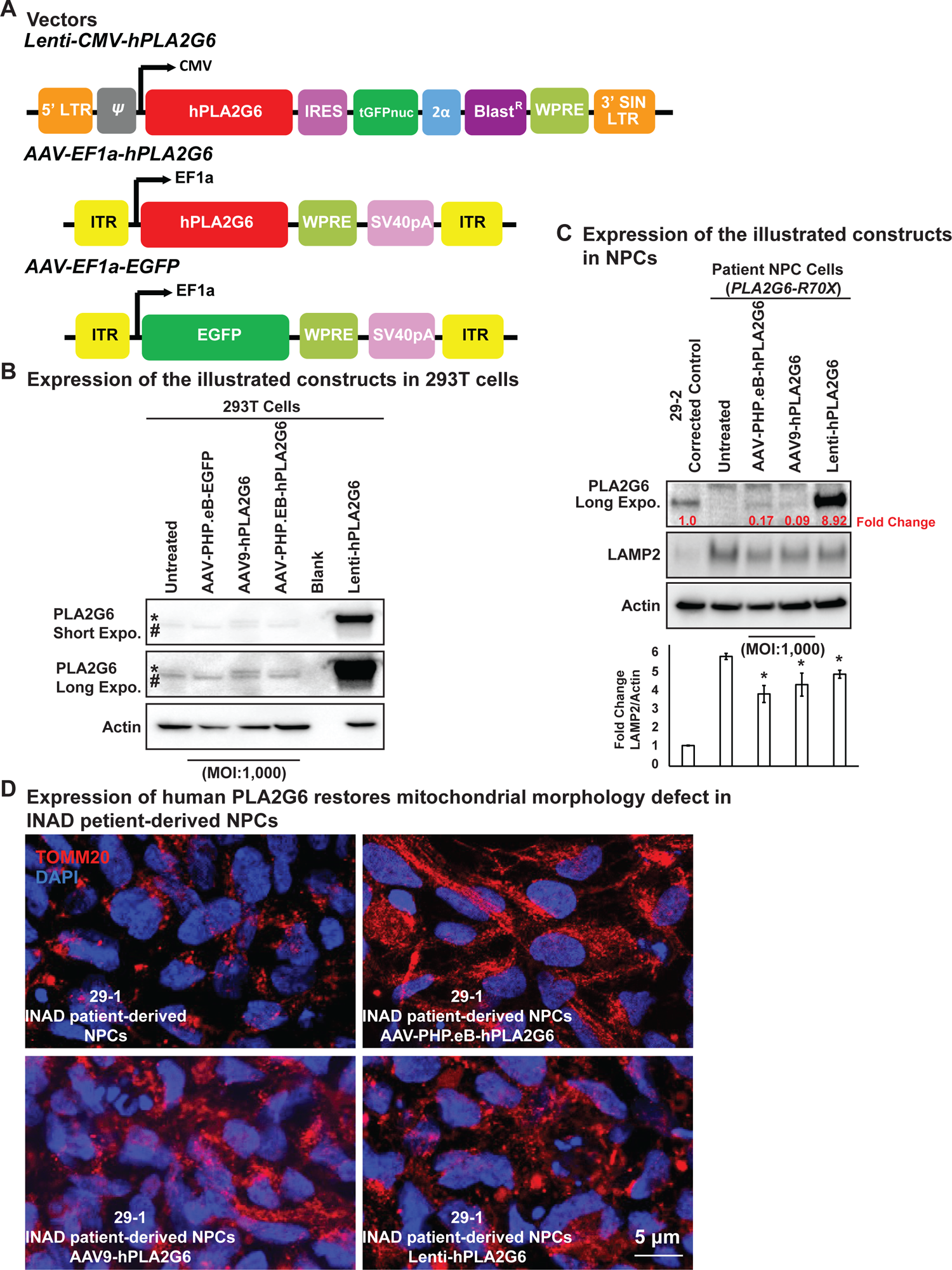
Expression of human PLA2G6 restores lysosomal and mitochondrial morphology defects in INAD patient-derived NPC lines. A. Vectors/constructs. B. Expression levels of the constructs (A) in 293T cells. PLA2G6 antibody was used to detect the endogenous PLA2G6 of cellular lysates. * represents endogenous PLA2G6. # indicates a non-specific band. All AAV constructs were used to infect 293T cell using a MOI of 1,000. The MOI of the Lenti-viral-based construct was not determined. C. Expression levels of the illustrated constructs (A) in NPCs. AAV constructs were used to infect NPCs at a MOI of 1,000. The MOI of the Lenti-viral-based construct was not determined. The intensity of LAMP2/Actin is quantified below (n=3). Error bars represent SEM; * P<0.05. D. Expression of human PLA2G6 restores mitochondrial morphology defects in INAD patient-derived NPCs.

### Pre-symptomatic injection of AAV-PHP.eB-hPLA2G6 suppresses rotarod defect and prolongs lifespan in KO/G373R INAD mice

To determine if delivery of *hPLA2G6* alleviates the phenotypes in INAD mouse model, we injected five *PLA2G6^KO/G373R^* mice with AAV-PHP.eB-EF1a-hPLA2G6 construct at postnatal day 40 via ICV and IV injections (Figure 5A; blue line). In an independent litter, we injected two more *PLA2G6^KO/G373R^* mice at postnatal day 40 via ICV only (Figure 5A; purple line). Un-injected *PLA2G6^KO/G373R^* mice displayed the first signs of rotarod impairment at ∼ 90 days of age (Figure 5A; red line). In contrast, the ICV and IV injected *PLA2G6^KO/G373R^* mice (Figure 5A; blue line) did not show signs of rotarod impairment until 130-140 days, a 50-60 day delay in the onset of the rotarod defects. However, even though the ICV-injected mice show some improvement in rotarod performance, statistical analyses did not show significance at most time points (Figure 5A; purple line). Note that injection of the AAV-PHP.eB-EF1a-PLA2G6 construct in wild-type littermates did not cause any obvious change in rotarod performance.

**Figure 5:**
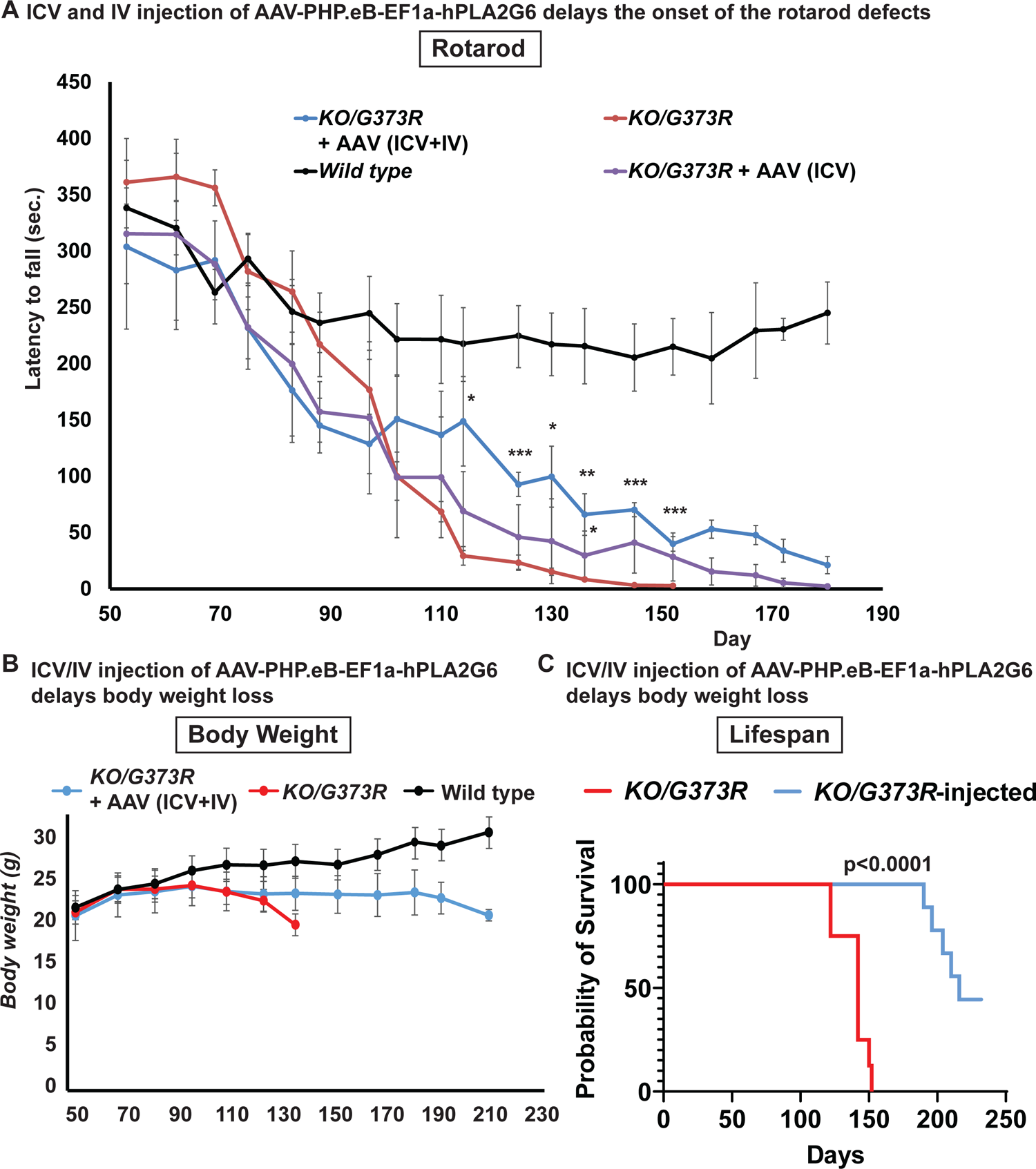
Pre-symptomatic injection of AAV-PHP.eB-hPLA2G6 suppresses rotarod defect and prolongs lifespan in *PLA2G6^KO/G373R^* INAD mice. A. Rotarod defect in *PLA2G6^KO/G373R^* mice (genotype: *PLA2G6^KO/G373R^*) (n=6) are reduced by pre-symptomatic (P40) ICV + IV injection (n=5), but not ICV only (n=2). Wild-type (n=5); *PLA2G6^KO/G373R^* (n=10). Rotarod performance of mice with the indicated genotypes was measured weekly. B. Pre-symptomatic (P40) ICV + IV injection of AAV-PHP.eB-hPLA2G6 stabilizes body weight of *PLA2G6^KO/G373R^* mice. Wild-type (n=5); *PLA2G6^KO/G373R^* (n=6); *PLA2G6^KO/G373R^* injected (n=3). C. Pre-symptomatic (P40) ICV + IV injection of AAV-PHP.eB-hPLA2G6 prolongs lifespan of the *PLA2G6^KO/G373R^* mice. *PLA2G6^KO/G373R^* (n=8); *PLA2G6^KO/G373R^* injected (n=8). Error bars represent SEM. * P<0.05; ** P<0.01; *** P<0.001. All assays were conducted in blind of genotypes and treatments.

We also assessed body weight and lifespan. All injected *PLA2G6^KO/G373R^* mice (both the ICV+IV injected group and IV injected group) exhibit a sudden body weight drop at ∼190 days, two weeks before they die (Figure 5B). This is ∼70 days later than the uninjected *PLA2G6^KO/G373R^* animals, which show an obvious drop in body weight at ∼120 days (Figure 5B). Half of the injected mice died by 210 days and four animals are still alive (Figure 5C; blue line). Hence, the injected animals live an average of at least 65 days longer than the uninjected *PLA2G6^KO/G373R^* mice (Figure 5C; red line). In summary, our data suggest that 1) Expression of human PLA2G6 in mice is safe; 2) Expression of human PLA2G6 in adult pre-symptomatic *PLA2G6^KO/G373R^* mice delays the onset of defects; and 3) promoting broad expression of hPLA2G6 using ICV and IV injections is more effective than ICV delivery only.

## Discussion

We previously documented that flies lacking *iPLA2-VIA*, the fly homolog of *PLA2G6*, have a dysfunctional retromer complex, accumulate ceramides, exhibit an expansion of lysosomes, and develop aberrant mitochondria (Lin et al., 2018). We showed that PLA2G6 interacts with VPS26 and VPS35 and is required for proper retromer function to promote the recycling of proteins and lipids. Hence, loss of *PLA2G6* leads to a lysosomal expansion and disrupts ceramide metabolism. Drugs that decrease ceramide levels have a beneficial effect and significantly improve the above phenotypes, arguing that ceramides play an important role in the pathogenesis. Accumulated ceramides stiffen membranes, cause mitochondrial defects, and contribute to a negative feed-forward amplification of the defects (Lin et al., 2018; Lin et al., 2019). Here, we turn to human cells and mice to establish if the same pathway is affected. We show that *PLA2G6* is highly expressed in iPSCs and NPCs but not in fibroblasts (Figure 1A). INAD patient-derived NPC and DA neurons that lack PLA2G6 also exhibit an accumulation of ceramides, expansion of lysosomes, and disruption of mitochondrial morphology (Figure 1 and Supplemental Figure 3). Similarly, we observed the same defects in two INAD mouse models (Figure 2 and Supplemental Figure 4). Our data indicate that these defects are a root cause of INAD/PARK14.

Cerebellar atrophy is one of the earliest features shared by most INAD patients based on MRI studies (Farina et al., 1999). In *PLA2G6* knockout mice, cerebellar atrophy, as well as a loss of Purkinje neurons were observed in older animals (18 months) (Zhao et al., 2011). Moreover, mice with a knock-in of a variant identified in *PARK14* patients, exhibit an early-onset loss of the substantia nigra and a loss of DA neurons (Chiu et al., 2019). These data suggest that loss of Purkinje neurons in the cerebellum and/or DA neurons in the substantia nigra are features that are shared by patients and mice that lack *PLA2G6*. Here, we show that GlcCer is highly enriched in Purkinje and DA neurons in mutant *PLA2G6* mice (Figure 2D) and in INAD patient-derived DA neurons (Figure 1B and Supplement Figure 3B). We also observed significant expansion of the lysosomes as well as mitochondrial defects in the mice DA neurons and Purkinje cells as well as in DA neurons derived from NPCs (Figure 1C-F, Figure 2C and Supplement Figure 4), consistent with the observed lesions in patients.

In the past two years, *PLA2G6* has been shown to function as a key regulator of ferroptosis in cancerous cell lines and in placental trophoblasts. Elevated levels of reactive oxygen species (ROS) and iron lead to ROS-induced lipid peroxidation. This promotes ferroptosis (Jiang et al., 2021). Loss of *PLA2G6* in cancerous cell lines or placental trophoblasts promotes lipid peroxidation and ferroptosis (Beharier et al., 2020; Chen et al., 2021; Kajiwara et al., 2022; Wang et al., 2022b). An accumulation of peroxidated lipids at day 25 was also observed in adult brains of flies that carry a hypomorphic allele of *iPLA2-VIA* (Kinghorn et al., 2015). Moreover, *PLA2G6^R748W/R748W^* mice that contain a PARK14 variant, exhibit an early impairment of rotarod performance and an elevation of ferroptotic death and a loss of DA neurons in the midbrains was observed in 7-month-old *PLA2G6^R748W/R748W^* knock-in mice (Sun et al., 2021). Interestingly, ferroptosis is also observed in rotenone-infused rats as well as in α-synuclein-mutant *Snca^A53T^* mice, suggesting that this pathway may be affected in different models of PD (Sun et al., 2021). However, in flies that lack PLA2G6, we did not observe an elevation of iron or ROS in 15-day-old adults (flies live a maximum of 30 days), yet these flies already exhibit a severe ceramide accumulation and lysosomal expansion. Similarly, the INAD mice exhibit a severe ceramide accumulation and lysosome expansion in Purkinje neurons and DA neurons prior to neuronal death. We, therefore, argue that ceramide accumulation and lysosome expansion precede ferroptosis. Our data are also consistent with the observation that activation of lysosomes and autophagy promote ferroptosis (Hou et al., 2016; Torii et al., 2016). Hence, loss of *PLA2G6* may disrupt retromer function, and lead to impaired recycling of proteins and lipids which in turn causes lysosomal and autophagy defects that activate ferroptosis.

We screened drugs that affect sphingolipid metabolism, endo-lysosomal trafficking as well as drugs that are being explored in PD. Upon screening these drugs in flies, we identified drugs that had a positive impact on bang-sensitivity and screened them in NPCs derived from an INAD patient. We identified four drugs, Ambroxol, Desipramine, Azoramide, and Genistein that reduce LAMP2 levels in INAD patient-derived NPCs (Figure 3). Upon oral uptake, Ambroxol is transported to lysosomes where it serves as a molecular chaperone for β-glucocerebrosidase (Magalhaes et al., 2018), the enzyme encoded by *GBA,* associated with Gaucher Disease and PD (Sidransky, 2004; Wong et al., 2004). Ambroxol promotes β-glucocerebrosidase activity in the lysosomes to reduce the levels of GlcCer (Magalhaes et al., 2018). Given that loss of *PLA2G6* leads to a robust accumulation of GlcCer in Purkinje neurons and in DA neurons, Ambroxol may promote the degradation of GlcCer and reduce the toxicity of GlcCer accumulation, consistent with our model and observations. However, it should not affect the elevation of other ceramides and hence may only partially suppress the phenotypes associated with elevated ceramides. Indeed, RNAi knockdown of *lace*, the rate-limiting enzyme that synthesizes ceramides, strongly impairs ceramide synthesis, and has the most potent suppressive effect as it affects all ceramides (Lin et al., 2018).

Interestingly, Miglustat and Ibiglustat, two inhibitors of UGCG UDP-glucose ceramide glucosyltransferase that are being used to reduce GlcCer levels in Gaucher disease patients, did not reduce the bang-sensitivity in INAD flies (Figure 3B). UGCG UDP-glucose ceramide glucosyltransferase is the enzyme that generates GlcCer. Blocking its activity reduces GlcCer levels but does not affect the other ceramides. We argue that this may lead to an elevation of other ceramides, and these drugs may not be effective given that many other ceramides are elevated in INAD. Indeed, Desipramine, a tricyclic antidepressant that is transported to the lysosomes where it functions as an acidic sphingomyelinase inhibitor to suppress the overall ceramide synthesis (Jenkins et al., 2011), showed a significant rescue effect (Figure 3). Hence, based on our drug screen, we identify two drugs, Ambroxol and Desipramine that reduce the levels of ceramides and lysosomal defects (Lin et al., 2018). These data are consistent with our model that increased ceramide levels contribute to the pathogenesis of INAD and *PARK14*.

Azoramide, a drug tested for Parkinsonism, promotes protein folding and secretion without inducing ER stress in the endoplasmic reticulum (Fu et al., 2015). It reduces ER stress, abnormal calcium homeostasis, mitochondrial dysfunction, elevated ROS as well as cell death in PARK14 patient-derived DA neurons (Ke et al., 2020). Azoramide reduces bang-sensitivity in flies and reduces the lysosomal expansion in INAD patient-derived NPCs (Figure 3). Azoramide promotes protein folding and hence may reduce the levels of misfolded proteins and alleviate lysosomal stress (Jackson and Hewitt, 2016). Finally, Genistein is an isoflavone naturally found in soy products. It exhibits neuroprotective effects in DA neurons in the MPTP-induced mouse model of Parkinson disease (Liu et al., 2008). It has been shown to promote lysosomal biogenesis by activating the transcription factor EB (TFEB) (Moskot et al., 2014). Taken together, the identification of Azoramide and Genistein to alleviate neurodegeneration in INAD models indicates that lysosomal defects are key contributors to the pathogenesis of INAD.

We previously showed that whole body expression of human *PLA2G6* cDNA in flies that lack *iPLA2-VIA* completely rescued the neurodegenerative defects as well as lifespan, whereas neuronal expression strongly suppressed the neuronal phenotypes but did not extend lifespan (Lin et al., 2018). To assess the effect of expressing PLA2G6 in INAD mice and patient NPCs, we created two AAV-based vectors, *AAV-EF1a-EGFP* and *AAV-EF1a-hPLA2G6*, which express EGFP or human PLA2G6 under the control of a ubiquitous promoter (*EF1a*) in human NPCs and mice. After delivery using ICV and IV into mice, the EGFP is broadly expressed in the nervous system as well as in other organs including the heart and liver (Supplement Figure 5). Upon delivery of *AAV-EF1a-hPLA2G6* to patient-derived NPCs, the construct expresses low levels of hPLA2G6 (10-20% of the endogenous levels). However, this is sufficient to partially alleviate the defects including lysosomal expansion and mitochondrial morphological abnormalities in NPCs (Figure 4). Moreover, delivery of *AAV-EF1a-hPLA2G6* into *PLA2G6^KO/G373R^* mice also delays the onset of rotarod defect, helps to sustain body weight, and prolongs lifespan (Figure 5). These proof-of-concept data are important because they indicate that the defects caused by a loss of *PLA2G6* can be delayed by low levels of *PLA2G6* expression and reversed in NPCs. However, the efficiency of delivery and expression need to be improved, possibly by testing other serotypes of AAV or injecting more virus.

In summary, the accumulation of ceramides, expansion of lysosomes and disruption of mitochondria are key phenotypes that are associated with the loss of *PLA2G6* in flies, mice, and human cells. The defects are obvious in mouse DA neurons and Purkinje cells, two cell populations that have been implicated in INAD/PARK14. We also identified four drugs that suppress the ceramide levels or promote lysosome functions and that alleviate the defects caused by loss of *PLA2G6* in INAD flies and in INAD patient-derived NPCs. These data, combined with previous data, provide compelling evidence that ceramides, expansion of lysosomes, and disruption of mitochondria are at the root of the pathogenesis of INAD and PARK14. Finally, we report encouraging data for a proof-of-concept trial to test the efficiency of a gene therapy approach. We argue that combining a drug and gene therapy approach will provide an avenue to significantly improve the quality of life of INAD/PARK14 patients.

## List of Supplementary Materials

**Figure 1-Figure Supplement 1:**
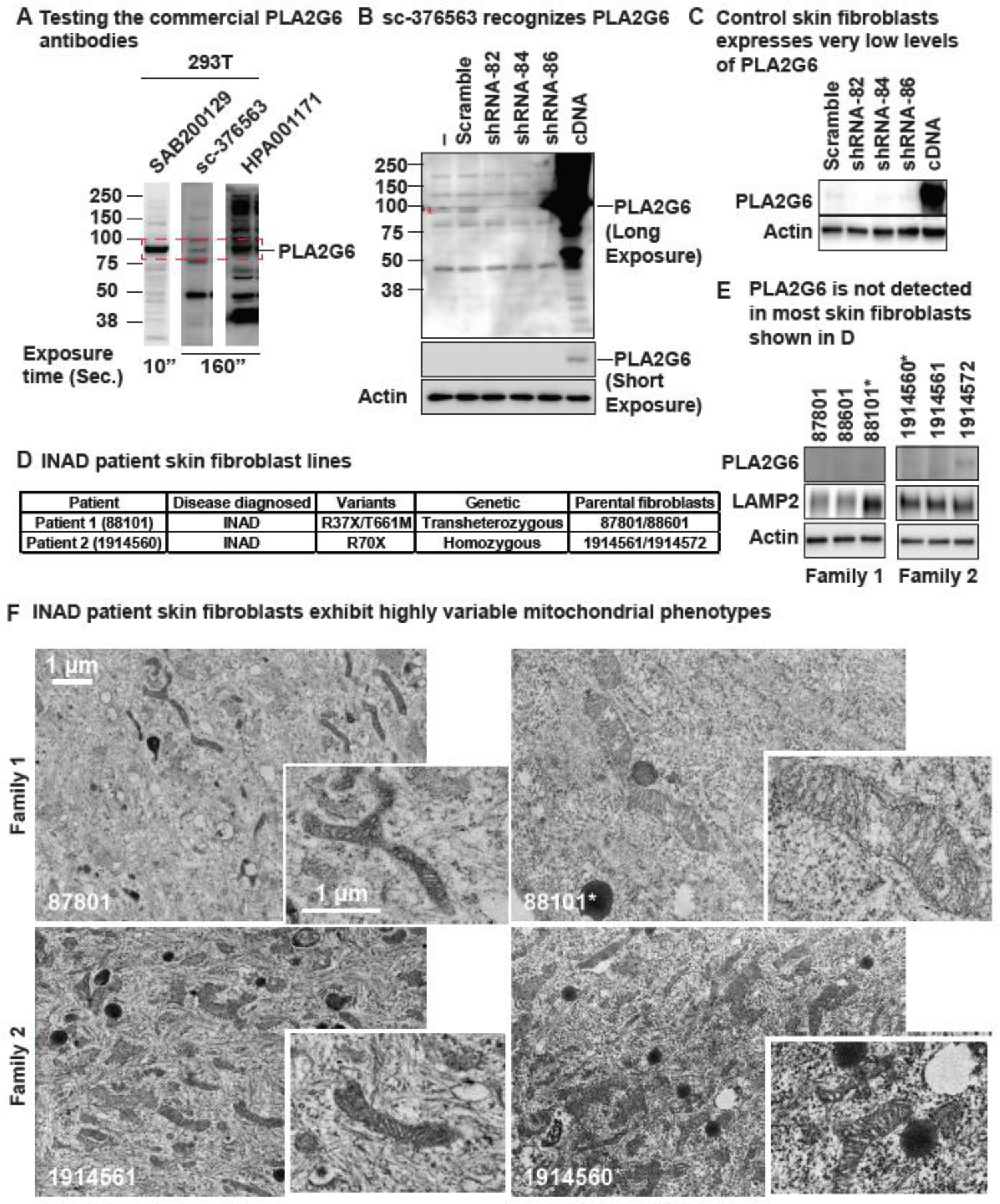
Human skin fibroblasts express no or very low levels of PLA2G6 and exhibit highly variable phenotypes. A. identification of commercially available PLA2G6 antibodies that specifically recognize endogenous levels of PLA2G6 in HEK293T cellular lysate. The SAB200129 antibody recognizes a major band at the predicted molecular weight (85/88 kDa) of PLA2G6. The sc-376563 antibody detects five major bands with one band at ∼85/88 kDa. HPA001171 detects too many bands at similar intensity, hence is not a proper antibody to be used. B-C. sc-376563 specifically recognizes endogenous PLA2G6 in HEK293T cells (B) and a control skin fibroblast line (C). The 85/88 kDa (the predicted molecular weight of PLA2G6) band detected by sc-376563 is significantly reduced upon PLA2G6 shRNA treatments, whereas the single band detected by SAB200129 does not. D. INAD patient-derived skin fibroblasts and the parental control fibroblasts. E. PLA2G6 is not expressed in most skin fibroblasts shown in D. The experiments were conducted in blind of the genotype. sc-376563 antibody was used to label PLA2G6. LAMP2 antibody was used to assess lysosomal accumulation. Actin was used as a loading control. F. INAD patient skin fibroblasts exhibit highly variable mitochondrial phenotypes. Scale bar = 1 µm.

**Figure 1-Figure Supplement 2.**
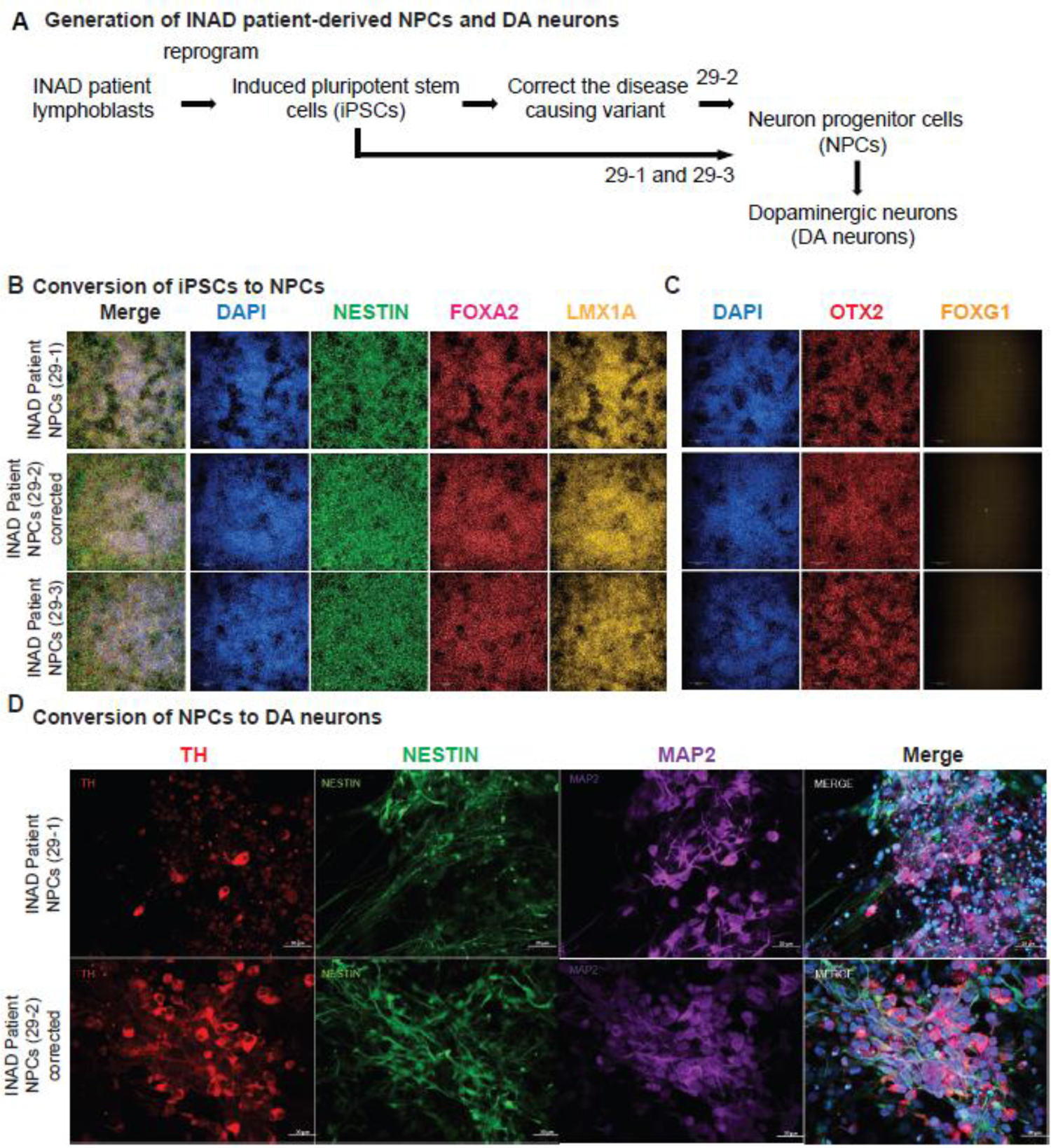
A. Generation of INAD patient-derived NPCs and DA neurons. B-D. Patient and isogenic control lines differentiate into comparable ventral midbrain floor plate neuron progenitor cells. B-C. Day 22 cultures from an INAD patient derived PLA2G6 mutant iPSC clone 29-1, isogenic PLA2G6 corrected iPSC clone 29-2 (derived from 29-1), and INAD patient derived PLA2G6 mutant iPSC clone 29-3. B. uniformly express pan NPC intermediate filament marker NESTIN (green), and nuclear (DAPI, blue), floor plate markers FOXA2 (red), and vM FP marker LMX1A. C. Sister wells show nuclei (DAPI, blue), uniformly express rostral-to-midbrain marker OTX2 (red) and do not express forebrain marker FOXG1 (yellow). 3×3 montage of representative randomly sampled 10x Phenix confocal fields. D. The differentiated DA neurons are TH (red), NESTIN (green) and MAP2 (purple) positive. Scale bars = 20 µm.

**Figure 1-Figure Supplement 3:**
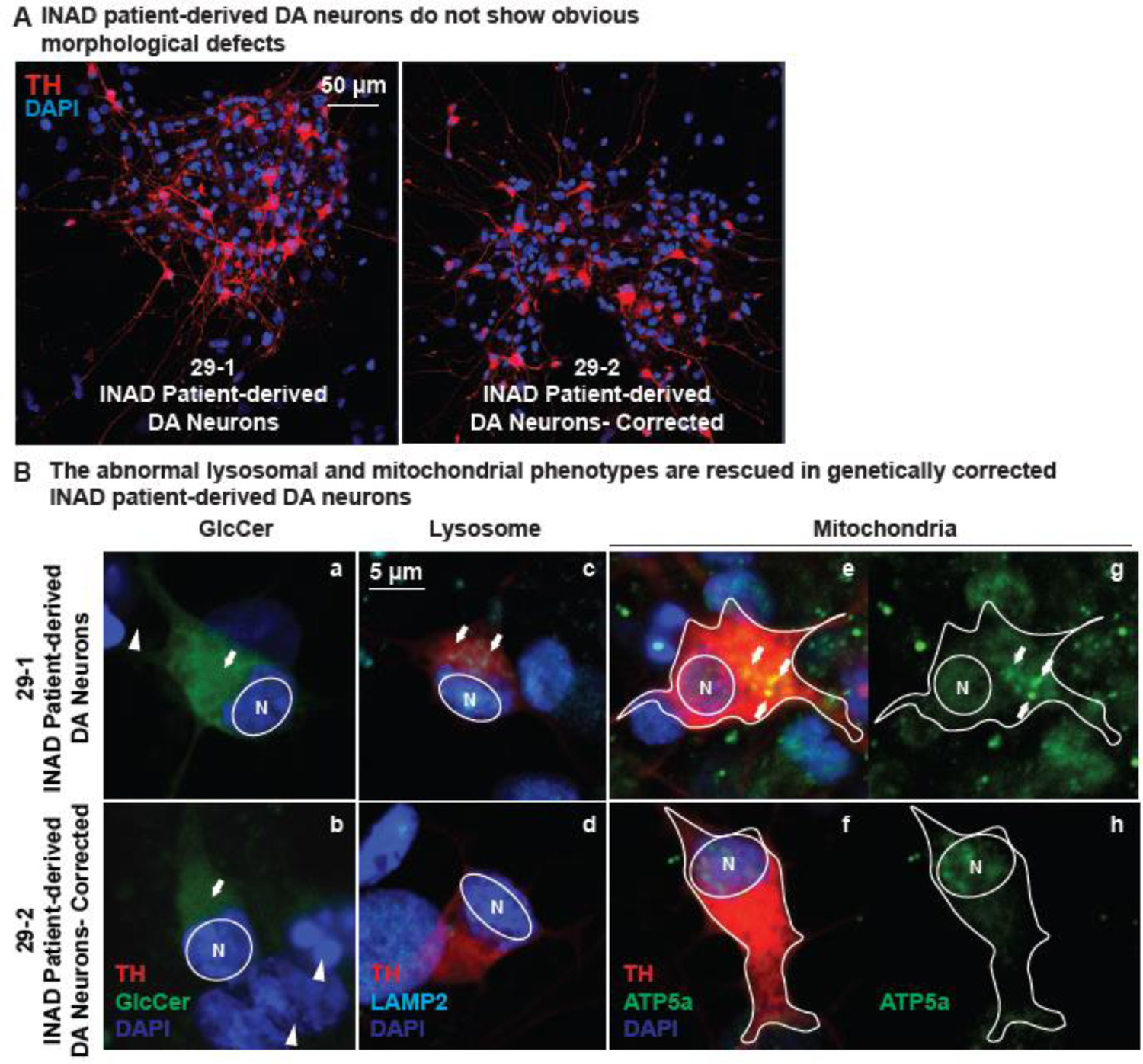
Ceramide accumulation, lysosomal expansion and mitochondrial defects in INAD patient-derived DA neurons. A. INAD patient-derived DA neurons do not show obvious morphological defects. TH (Tyrosine Hydroxylase, red) antibody labels DA neurons. DAPI (blue) labels cell nuclei. Scale bar = 50 µm. B. The abnormal lysosomal and mitochondrial phenotypes are rescued in genetically corrected INAD patient-derived DA neurons. GlcCer antibody (green; arrow in a and b), LAMP2 antibody (Cyan; arrows in c and d), and ATP5a antibody (green; arrows in e-h) were used to label GlcCer, lysosomes and mitochondria, respectively. Arrowheads indicate the undifferentiated NPCs. Scale bar = 5 µm.

**Figure 2-Figure Supplement 1:**
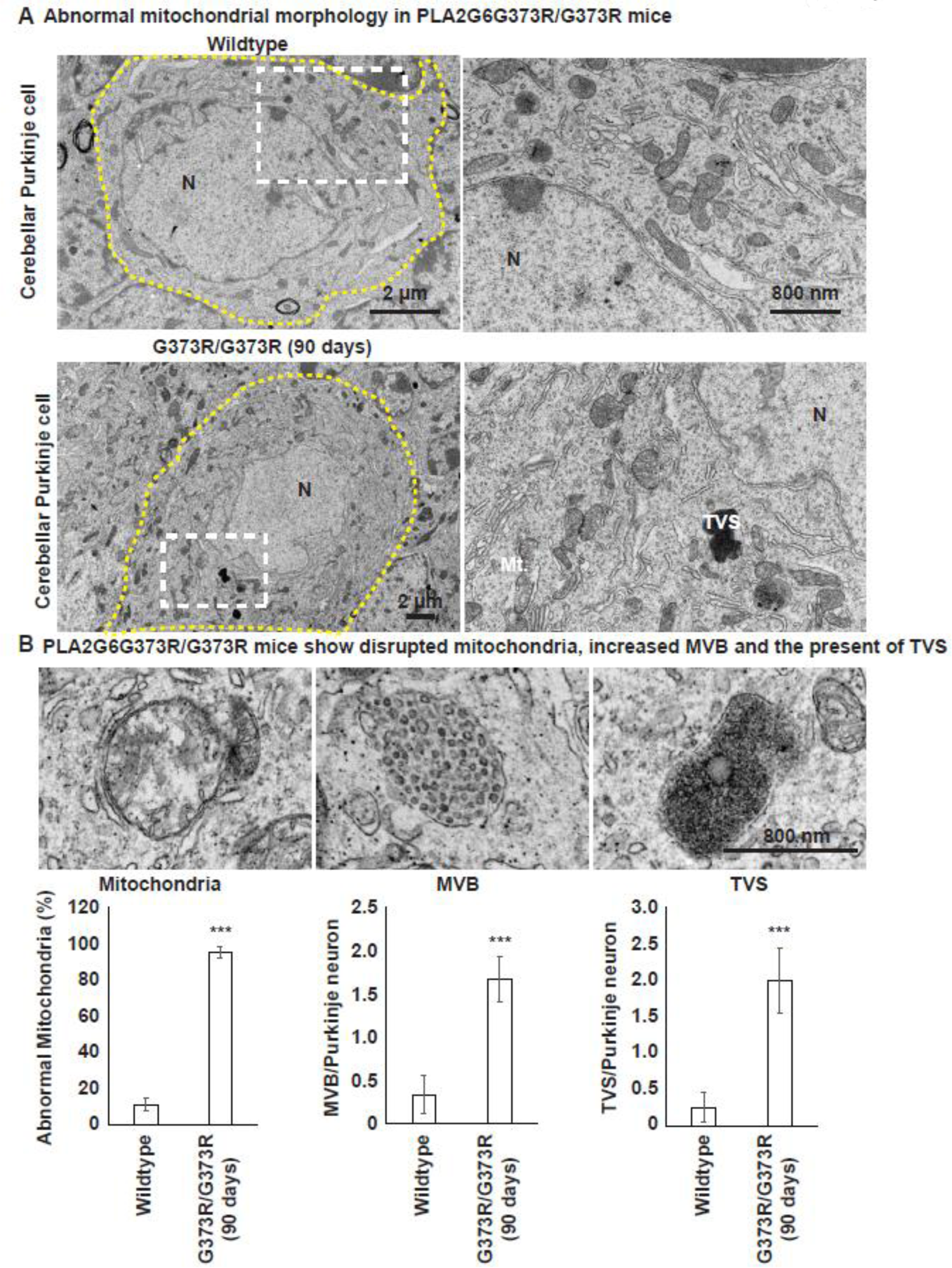
*PLA2G6^G373R/G373R^* mice show disrupted mitochondria, increased MVB and the present of TVS in Purkinje neurons. A. Abnormal mitochondrial morphology in Purkinje neurons of *PLA2G6^G373R/G373R^* mice. The yellow dotted line circles a representative Purkinje neuron. Scale bar = 2 µm. The boxed regions are enlarged at right. Scale bar = 800 nm. Mt: mitochondria; N: nuclei; TVS: tubulovesicular structure. B. *PLA2G6^G373R/G373R^* mice show disrupted mitochondria, increased MVB and the present of TVS. The quantification of the disrupted mitochondria, number of MVB or TVS in the Purkinje neurons of *PLA2G6^G373R/G373R^* mice. Error bars represent SEM. *** P<0.001.

**Figure 5-Figure Supplement 5:**
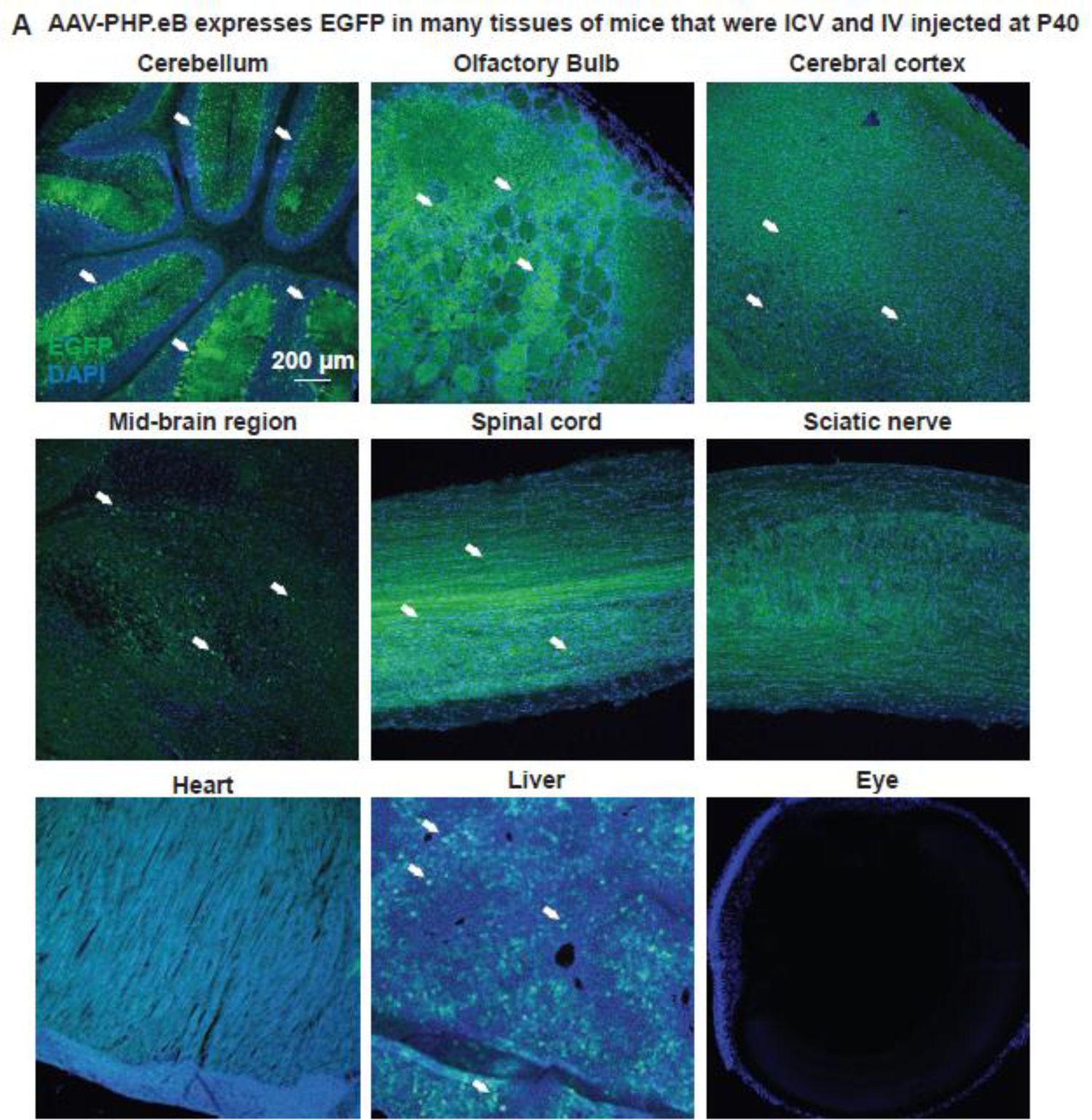
*AAV-EF1a-EGFP* injected via ICV and IV at P40 expresses EGFP in the indicated sites/tissues of wild type mice. GFP antibody was used to detect EGFP in the indicated sites/tissues. Arrows indicate cells that express high levels of EGFP. Scale bar = 200 µm.

## Acknowledgments

We thank Drs. Jin Xu for very insightful comments. We thank the Gene Vector Core at Baylor College of Medicine for packaging the AAV-PHP.eB virus. We thank the GERM Core at Baylor College of Medicine for the *in vitro* fertilization to retrieve the *PLA2G6^G373R^* mice. We are grateful for the generous gift of skin fibroblasts from Dr. Bénédicte Heron (Neurologie Pédiatrique Hôpital Trousseau) and Dr. Young-Hui Jiang (Duke University). This project was supported in part by Baylor College of Medicine IDDRC Grant Number P50HD103555 from the Eunice Kennedy Shriver National Institute of Child Health and Human Development for use of the Microscopy Core facilities. We acknowledge the Shan and Lee-Jun Wong Fellowship to GL. We thank the INADcure Foundation for support to GL. HJB and the NYSCF. This project was also supported by the Huffington Foundation and the Neurological Research Institute of TCH to HJB.

## Author contribution

GL: conception and design, acquisition of data, analysis and interpretation of data, drafting and revising the article. BT: conception and design, analysis and interpretation of data, and revising the article. GM, RCT and GC: acquisition of data, analysis and interpretation of data, and revising the article. LP and AH: analysis and interpretation of data, and revising the article. AJHL, ZZ, and LW: acquisition of data. HJB: conception and design, analysis and interpretation of data, and drafting and revising the article.

## Materials and Methods

### *Drosophila* and drug treatment

Mixed genders of flies (approximately 50 % male and 50 % female) were used for all experiments. Flies were raised on molasses-based food at 25 °C in constant darkness. The genotypes of the flies used is *y w;; iPLA2-VIA^Δ174^* (Lin et al., 2018). All drugs were added freshly to regular fly food at the indicated concentration. The flies were transferred to fresh food with or without the drugs every three days.

### *Drosophila* behavioral assay

To perform bang-sensitive paralytic assays, five adult flies were tested per vial. The flies were vortexed at maximum speed for 15 sec and the time required for flies to stand on their feet was counted. At least 60 flies were tested per data point.

### Fibroblasts from INAD patients and their parents

Human skin fibroblasts from Family 1 and 2 were a gift from Dr. Bénédicte Heron (Neurologie Pédiatrique Hôpital Trousseau) and Dr. Young-Hui Jiang (Duke University), respectively. The control skin fibroblasts (cat. #GM23815) were purchased from Coriell Institute.

### Human iPSC (hiPSC) culture

hiPSCs maintained on Cultrex (CTX Cat# 3434-010-02, R&D Systems) in PSC Freedom Media (FRD1, ThermoFisher Custom) were passaged every 4-5 days using Stem-Accutase (Cat# A11105-01, Life Technologies) in the presence of 1 µM Thiazovivin (THZ Cat# SML1045-25 MG, Sigma-Aldrich).

### CRISPR/Cas9-mediated gene editing

#### Transfection

4 x10^5^ Stem-Accutase dissociated hiPSCs were plated onto a CTX pre-coated 24 well in FRD1 containing THZ. Cells were transfected directly after passaging with the transfection reaction. A RNP complex was formed by incubating 18 pM Alt-R® S.p. Cas9 Nuclease V3 (Cat# 1081058 IDT Technologies) with 18 pM PLA2G6 Alt-R® CRISPR-Cas9 sgRNA (GTGAGTTCCTGGGGTTGACC, IDT) for 5 min at RT. The RNP complex was then incubated for 15-20 min at RT with 120 uM Alt-R HDR Oligos (tccaatccgagacgtgggggagtgaaaggagagaagtatgttcccgctgagcatcacccaccggaatccactctgtgagttcctggggttgaccaAgacgcagtcccaggtgcggttgggagtgttc, IDT Technologies), 2 µl Lipofectamine Stem (ThermoFisher, STEM00008,) and 40 µl Opti-MEM (Cat# 31985062, ThermoFisher).

#### Monoclonalization

Transfected iPSCs were single cell sorted into 96 well plates using a Benchtop Microfluidic Cell Sorter (Nanocollect). Plates were fed daily with FRD1 and scanned every night on a Celigo Image Cytometer (Nexcelom Bioscience). After 10 days monoclonal colonies were consolidated and passaged into a new 96 well plate. Wells were passaged when reaching 80-100% confluency for freeze backs and sequencing analysis.

#### Sanger sequencing of monoclonal wells

30 µl of Quick Extract DNA Estraction (Lucigen, QE09050,) was added to 5.0 x 10^4^ pelleted iPSC, resuspended and incubated for 15 min at 65 ⁰C. Quick extract lysate template was prepared by positing 5.0 x 10^4^ cells into a 96-well hard-shell PCR plate (Bio-Rad). The PCR for sanger sequencing was performed by using 2 µl of quick extracted gDNA in a 25 µl PCR reaction using AmpliTaq Gold 360 and PLA2G60 primer pairs (fwd: gccgcctggtcaataccttc, rev: acccctcagacagagactcaa). The amplicon was send for Sanger sequencing subsequently.

#### Quality Control Measures

iPSCs were expanded via automation on the NYSCF Global Stem Cell Array platform for further quality control assays and then frozen into barcoded Matrix tubes in Synth-a-freeze Cryopreservation Media at R500k cells/vial.

All iPSC lines undergo rigorous quality control that includes a sterility check, mycoplasma testing, viability, karyotyping via Illumina Global Screening Array, SNP ID fingerprinting via Fluidigm SNPTrace, pluripotency and embryoid body scorecard assays via Nanostring. iPSCs were maintained using Freedom (ThermoFisher, custom) media.

### Generation of the isogenic INAD patient-derived iPSC, NPCs and DA neurons

Undifferentiated iPSCs were grown following standard protocols in StemFlex medium on laminin521 substrate (BioLamina) as described previously (Ruzo et al., 2018). iPSCs were differentiated to ventral midbrain neural progenitor cells (NPC) and dopaminergic (DA) neurons following (Kim et al., 2021) with the following modifications. Neural induction was initiated at day 0 by passaging iPSC, single cell seeding at 1M cells/ml in 2.5M/well of over ultralow adherence plastic 6wp (Corning, cat#3471vendor) in neural induction medium (base medium changed to AaDMEMEMM/F12:Neurobasalbobasla 50:50, with N2, B27 without RA, RhoK inhibitor Y27632 10uM (first 2 days), 100 nM LDNn, 10 uM SB-431542 and, substituting 1 mM SAG3.1 and 1 mM Prurmorprphhpamione (R&D) in place of SHH protein,) and 1uM CHIR99021 (R&D). At day 4-7 CHIR99021 was increased to 6 µM, and from day 8-11 was decreased to 3 µM and 0.2 mM AA,(Sigma-Aldrich), 0.2 mM dbcamp (Sigma-Aldrich), 10 µM DAPT (Tocris) and 1 ng/ml TGFb3 (R&D) 0 were added from day 10 onward. EBs were dissociated with accutase at day 16, and reseeded at 0.8Mcells/cm^2^ on polyornithine laminin (1 mg/ml in pH 8.4 borate buffer followed by 10 µg/ml natural mouse laminin in DMEM/F12) coated TC plastic for continued differentiation and day-22 fixation and staining for positional markers, or frozen in 2x FM (Millipore Sigma; ES-002-10F). Live cultures or thawed cells were either expanded as floor plate progenitors (following Brundin approach; Floor Plate Progenitor Kit expansion protocol (ThermoFisher; A3165801) or terminally differentiated to DA neurons with all prior factors except in neurobasal base medium after day 25.

### Western blotting

Cells were homogenized in 1 % NP40 lysis buffer (20 mM HEPES pH7.5, 150 mM NaCl, 1 % NP-40, 10 % Glycerol and Roche protease inhibitor mix) on ice. Tissue or cell debris were removed by centrifugation. Isolated lysates were loaded into 10 % gels, separated by SDS-PAGE, and transferred to nitrocellulose membranes (Bio-Rad). Primary antibodies used in this study were as follows: mouse anti-PLA2G6 antibody (Santa Cruz; sc376563), rabbit anti-PLA2G6 antibody (Sigma; SAB4200129); rabbit anti-PLA2G6 antibody (Sigma; HPA001171); mouse anti-Actin (ICN691001, ThermoFisher); and rabbit anti-LAMP2 antibody (Abcam; ab18528).

### Immunofluorescence staining

The cultured cells were fixed with 4 % paraformaldehyde in 1X PBS at 4 °C for 30 minutes. The fixed cells were permeabilized in 0.1 % Triton X-100 in 1X PBS for at least 15 minutes at room temperature. For mouse tissues, the mouse were deeply anesthetized using isoflurane, and perfused intracardially with 1X PBS and 4 % paraformaldehyde. Brains were dissected and post fixed in 4 % PFA/PBS overnight at 4°C. The next day, tissues were cryoprotected in a 20 % sucrose/PBS solution at 4 °C for one day, followed by a 30% sucrose/PBS solution at 4 °C for one more day. Tissues were then embedded and frozen in OCT and cryosectioned using a cryostat (Leica CM1860) at 25–40 μm. The fixed mouse tissues were permeabilized in 0.1 % Triton X-100 in 1X PBS for at least 15 minutes at room temperature. The following antibodies were used for the immunofluorescence staining: Chicken anti-Tyrosine Hydroxylase antibody (Abcam; ab76442; RRID:AB_1524535); rabbit anti-LAMP2 antibody (Abcam; ab18528; RRID:AB_775981); mouse anti-TOMM20 antibody (Abcam; ab56783; RRID:AB_945896); Rabbit anti-GlcCer (Glycobiotech; RAS_0010); Mouse anti-ATP5a (Abcam; ab14748; RRID:AB_301447); mouse anti-LAMP1 (Abcam; ab25630; RRID:AB_470708); mouse anti-GFP FITC conjugated (Santa Cruz; sc9996; RRID:AB_627695); NESTIN (Millipore; ABD69; RRID:AB_2744681); FOXA2 (R & D; AF2400; RRID:AB_2294104); LMX1A (Millipore; AB10533; RRID:AB_10805970); OTX2 (R & D; AF1979; RRID:AB_2157172), FOXG1 (Takara; ABM227), and Alexa 488-, Cy3-, or Cy5-conjugated secondary antibodies (111-545-144, 111-585-003 and 111-175-144, Jackson ImmunoResearch Labs; or Alexa 488, 555, 647 conjugated anti-rabbit, mouse, and goat (ThermoFisher). All the confocal images were acquired with a Model SP8 confocal microscope (Leica), with the exception of vmNPC characterization and initial Da neuron characterization, acquired with a Phenix spinning disk confocal HTS microscope (PE) and LSM700 Confocal microscope (Zeiss) respectively. Confocal images were processed using Image J and Photoshop (Adobe).

### Molecular cloning of the gene therapy constructs

Construction of pAAV-EF1a-hPLA2G6: PCR amplified fragments of the full length human PLA2G6 cDNA (NM_001349864.2) containing XbaI (5’) and EcoRV (3’) were digested with XbaI and EcoRV. To generate the vector backbone, pAAV-EF1a-CVS-G-WPRE-pGHpA (Addgene; 67528) was digested with XbaI and EcoRV and gel eluted to remove the CVS-G. The digested PCR fragments were ligated in the digested vector backbone to generate pAAV-EF1a-hPLA2G6-WPRE-pGHpA. The pGHpA was replaced by SV40pA to reduce the total size of the construct. Similarly, the pAAV-EF1a-EGFP was constructed following the same method except the EGFP cDNA was used. Lenti-CMV-hPLA2G6 was purchased from Horizon Discovery-Precision LentiORF collection (OHS5897-202617053).

### AAV-PHP.eB package

A plasmid DNA cocktail solution was produced by combining the pAAV transgene (pAAV-EF1a-hPLA2G6 or pAAV-EF1a-EGFP), rep/cap serotype PHP.eB, and AdΔF6 helper plasmids and transfected into HEK293T cells with iMFectin poly transfection reagent – GenDEPOT®. A total of 80 x 15-cm dishes were overlaid with the DNA cocktail solution and were allowed to incubate for 4-hours before adding fresh media. Three days post-transfection, dishes were imaged, harvested and digested. The collected cell viral lysate was centrifuged, supernatant was transferred to a new tube and digested separately. The cell pellet was resuspended in TMN (Tris-HCl cell suspension buffer) and cell associated AAV was recovered by cell lysis treatment with 5% sodium deoxycholate, DNase, RNase and 3 subsequent freeze/thaw cycles. Media secreted AAV was precipitated in a 40 % polyethylene glycol solution. Digest CVL was centrifuged to remove cell debris and cleared supernatant was transferred to a new tube. PEG precipitated supernatant was centrifuged, the resulting pellet was resuspended in HBS (HEPES Buffered Saline) solution. After digestion and precipitation, cleared cell viral lysate and media secreted AAV were combined and purified on a discontinuous iodixanol gradient. The band corresponding to purified AAV was extracted and diluted in DPBS supplemented with 0.001 % Pluronic F-68. Viral diluent was then concentrated in Amicon centrifugation filtration units (100,000 MW) to the desired volume. Concentrated AAV was first diluted 1:100 and then serially diluted 10-fold to yield AAV dilutions of 0.01, 0.001, 0.0001, and 0.0001. The titer of the AAV vectors were quantified with the primers corresponding to WPRE and probed against a GVC in-house standard. The AAV-PHP.eB was packaged by the Gene Vector Core at Baylor College of Medicine.

### Mouse house keeping

All experimental animals were treated in compliance with the United States Department of Health and Human Services and the Baylor College of Medicine IACUC guidelines. Mice were reared in 12-hour light-dark cycles with access to food and water ad libitum. The mouse lines used here include the *PLA2G6* complete knockout line (genotype: *PLA2G6^KO/KO^*) (Bao et al., 2004) and the *PLA2G6^G373R^* point mutation mouse line (Genotype: *PLA2G6^G373R/G373R^*) (Wada et al., 2009). The *PLA2G6* complete knockout line was a donation from Dr. Sasanka Ramanadham at the University of Alabama at Birmingham. Cryopreserved sperm of the *PLA2G6^G373R^* mice (No.BRRC04196) was purchased from RIKEN BioResource Research Center. The cryopreserved sperms were used to *in vitro* fertilize female C57BL/6 mice to retrieve the line. The *in vitro* fertilization was performed by the GERM Core at Baylor College of Medicine. After the *PLA2G6^G373R^* point mutation mice were retrieved, we mated *PLA2G6^KO/+^* with *PLA2G6^G373R/+^* mouse to generate transheterozygous mice (Genotype: *PLA2G6^KO/G373R^*). Mice were genotyped by standard PCR using isolated tail DNA as template.

### Mouse stereotaxic CNS injection

Preoperatively, adult mice were weighed and given meloxicam 30 minutes prior to surgery with a 30-gauge needle to minimize discomfort. Animals were anesthetized preoperatively with isoflurane. Following anesthesia, animals were checked for pedal withdrawal reflex; if absent, the animals were transferred to stereotaxic apparatus and maintained under anesthesia using volatilized isoflurane (1-3 % depending on physiological state of the animal, which was continuously monitored by response to tail/toe pinch). Isoflurane was diluted with pressurized oxygen using an anesthetic vaporizer system that allows precise adjustment of isoflurane and gas flow to the animal. Surgical site was prepared by shaving hair, administering depilatory cream and three times betadine surgical scrub, three times ethanol wipe. A short incision (.5-2 cm) was made over the skull using small surgical scissors. Following the incision, a small burr hole craniotomy (<1 mm in diameter) was made using a dental drill. 5 µl of AAV particles were injected into the third ventricle using a 33-gauge Hamilton syringe. Post-injection, the scalp was sutured using surgical nylon monofilament (Ethicon cat# 1689G) in a simple interrupted pattern for skin-to-skin closure. 7 days later, mice were briefly anesthetized with isoflurane and sutures were gently removed using small scissors and forceps.

### Mouse tail vein injections

Random selected mice were placed in restrainer to immobilize and have easy access to the tail. Tail was soaked in warm water for 10-15 seconds to cause vasodilation (enlargement) of the vein and was swabbed with a gauze dampened with alcohol to increase the visibility of the vein. One of the two lateral tail veins was located in the middle third of the tail. With the bevel of the needle facing upward and the needle almost parallel to the vein, the needle was slid into the tail vein. By gently applying negative pressure to the plunger and observing a flash of blood in the hub vein penetration was confirmed. Slowly the plunger was pressed to inject AAV-containing solution into the vein. The needle was removed from the vein and slight pressure was applied to the puncture site with a dry piece of gauze until the bleeding has stopped. The animal was removed from its restrainer and placed in the cage. The animal was monitored for 5-10 minutes to ensure hemostasis and that bleeding has stopped.

### Mouse rotarod assay

We measured the time (latency) a mouse takes to fall off the rod under continuous acceleration. On the day of testing, mice were kept in their home cages and acclimated to the testing room for at least 15 min (acclimation phase). The apparatus was set to accelerating mode from 5 to 40 rpm in 300 seconds. We record the latency at which each mouse falls off the rod. The first trial is the training period. During the training period, the mice were tested three times separated by 15 min intervals in a day and the trials were repeated for three consecutive days. After the training period, the mice were tested once a week. At the end of the rotarod assay, we weighted each mouse every other week. All the mice were tested in blind of genotype and treatments.

### Quantification and statistical analysis

For fly experiments, sample sizes are stated in the figure legends. The drug treatments were performed for 3 biological replicates. For all cell experiments, the studies were conducted in parallel with vehicle controls in the neighboring well for at least 3 biological replicates. All mouse experiments were conducted in the number of technical replicates indicated in the figure legends. Error bars are shown as standard error of the mean (SEM). The mouse survival datasets (Figure 2B and Figure 5C) were organized and analyzed in GraphPad Prism 9.4.0 using Mantel-Cox and Gehan-Breslow-Wilcoxon tests. All other datasets from flies and cells were organized and analyzed in Microsoft excel 2010 using Two-tailed Student’s t-test. The criteria for significance are NS: not significant; p>0.05; *p<0.05; **p<0.01 and ***p<0.001

